# Modeling glenohumeral stability in musculoskeletal simulations: A validation study with *in vivo* contact forces

**DOI:** 10.1101/2025.05.08.652806

**Authors:** Ibrahim Mohammed I. Hasan, Italo Belli, Ajay Seth, Elena M. Gutierrez-Farewik

## Abstract

Common optimization approaches to solve the muscle redundancy problem in musculoskeletal simulations can predict shoulder contact forces that violate joint stability with lines of action outside the glenohumeral joint border. Approaches with simple joint stability constraints were previously introduced imposing an upper limit on the direction of the contact force to stay within a specified stability perimeter. Such approaches predicted higher rotator cuff muscle activation than without constraints, but the estimated joint contact forces were oriented along the specified perimeter, raising questions about validity. In this study, several glenohumeral stability formulations were investigated, and tested against *in vivo* measurements of glenohumeral contact forces from the *Orthoload* dataset on one participant data in three dumbbell tasks: lateral raise, posterior raise, and and anterior raise. The investigated formulations either imposed inequality constraints on the contact force direction to remain within a stability perimeter whose shape was varied, or added a penalty term as a criterion measure to the objective function that made the objective function costly for contact force directions to deviate from the glenoid cavity center. All stability formulations predicted contact force magnitudes that agreed relatively well to the *in vivo* measured forces except for the strictest formulation that constrained the joint contact force to be directed at the glenoid cavity center. Models that restricted the force direction to lie within a specified shape estimated force vectors that largely lay along the perimeters. Models that instead penalized force directions that deviated from the glenoid cavity center estimated relatively more accurate contact force directions within the glenoid cavity, though still not entirely in agreement with *in vivo* measurements. Our findings support the proposed penalty formulations as more reasonable and accurate than other investigated existing glenohumeral stability formulations.

**Author summary:** In musculoskeletal models, the glenohumeral joint is often simplified as a purely rotational joint with no translation, whereas the actual joint movement involves both rotation and some translation, requiring stabilizing forces to prevent dislocation. Models that compute muscle forces based on a minimal effort strategy without specifically addressing glenohumeral stability may underestimate co-activation of the stabilizing rotator cuff muscles, and inaccurately predict that the contact force between the humerus and the glenoid is directed outside the articular surface of the joint. In this study, we compared existing stability models and proposed a new approach that penalized joint contact force whose direction deviates from the glenoid cavity center. We integrated these approaches into a muscle redundancy solver and estimated muscle and joint forces using the thoracoscapular shoulder musculoskeletal model. We compared model estimates to *in vivo* measurements of joint contact forces. We found that the penalty formulation reproduced the contact force direction most accurately, and promoted greater muscle co-contraction. These findings support the proposed penalty approach as a benchmark for more accurate analysis of shoulder biomechanics using musculoskeletal simulations.

## Introduction

The shoulder has the greatest range of motion in the human body, but is particularly prone to injuries in the glenohumeral (GH) joint [1, 2]. The GH joint is often considered as a ball-and-socket joint, but the socket, or the glenoid fossa of the scapula, is relatively shallow and small, covering at most only a third of the ball, or the head of the humerus [3]. Without surrounding bony cover to provide stability, GH stability is attained passively through glenoid concavity, the glenoid labrum surrounding the glenoid fossa, the joint capsule, ligaments and passive muscle tension, and actively through muscle contraction [4]. Any compromise in these stabilizing elements can lead to excessive humeral head translation and joint luxation, with increased risk of injury and pain [5]. Understanding joint stability is thus essential for preventing and mitigating shoulder injuries.

GH stability can be mechanically characterized by the magnitude and direction of the glenohumeral joint contact force (GH-JCF) vector. GH-JCF is the net sum of internal and external forces acting on the joint. The stability limit was introduced as a metric to quantify physical limits of GH stability [6, 7]. Stability limits were obtained experimentally via *in vitro* cadaver concavity compression tests wherein the humeral head was compressed into the glenoid cavity with a known force and shear force was applied perpendicularly until the humeral head was dislocated. The stability limit is defined as the magnitude ratio between the component of the GH-JCF that can translate the humeral head out of the glenoid cavity and the compressive component, which pushes the humeral head into the glenoid cavity. Higher stability limit in a given direction indicates a relatively more GH stability in that direction. If the stability limit is exceeded, the contact force vector will be directed outside the edge of the glenoid cavity, leading to joint dislocation. Empirical stability limits were reported as highest in the superior direction (≈ 0.60) and lowest in the anterior (≈ 0.32) direction with the glenoid labrum intact [7], indicating that GH joint is more stable in the superior-inferior direction than in the anterior-posterior direction.

Quantitative analysis of GH stability is contingent on the availability of either measurements or estimations of GH-JCFs. To quantify the GH-JCF, internal muscle and ligament forces in the joint must be known. Measuring activation of superficial muscles, e.g. deltoids, biceps and triceps, is easy to perform experimentally using surface electromoygraphy (EMG), however measuring activation of deep muscles, including rotator cuff muscles, would require extensive invasive fine-wire EMG measurements [8] which is experimentally difficult. Subsequent mapping of EMG measurements into muscle forces is furthermore still a major challenge [9].

Musculoskeletal modeling is an alternative approach to estimate muscle, ligament, and joint contact forces that are difficult to measure experimentally. Several shoulder musculoskeletal models exist and have been used to study shoulder biomechanics [10–17]. These models have been used to provide insights on shoulder loading in different applications, such as tendon transfer surgery [18], shoulder implants [19], and wheelchair propulsion [17, 20]. However, they have oversimplified the complex multibody geometry and articulations, specifically by ignoring [21] or inaccurately estimating the articulation of the scapula over the thorax [22, 23]. To address some of these shortcomings, a recently developed *OpenSim* shoulder model provided a novel formulation for scapulothoracic articulation that was validated with bone-pin kinematic measurements and shown to be robust for noisy marker tracking [24]. An enhanced version of this model, the thoracoscapular model, included several shoulder muscles such as the rotator cuff muscles and was used to study muscle work in typical shoulder movement [25], with muscle redundancy solved via the Computed Muscle Control algorithm [26].

In musculoskeletal simulations, the muscle redundancy problem has historically been addressed as an optimization problem that seeks a solution that satisfies a specified criterion measure and a set of imposed constraints. Frequently used criterion measures include minimum sum of muscle stresses [14, 27], muscle metabolic rates [27], or muscle activations [8, 13]. Unrealistic solutions to the muscle redundancy problem may arise if GH stability is not properly addressed [14, 22, 28, 29]. Unless restrained to stay within a defined stability perimeter, GH-JCFs estimated from musculoskeletal simulations could be directed outside of the articular surface of the glenoid accompanied by unrealistically low co-contraction of the rotator cuff muscles, though in reality this would suggest a tendency for instability and/or dislocation.

GH stability has been formulated as an inequality constraint that restrains the direction of the joint contact force to lie within a specified perimeter that describes the glenoid cavity, which has been estimated with different geometric shapes, including an ellipse [8, 10, 11], a circle [30] and a polynomial [28]. The circle model assumes that the GH stability is radially symmetric around the cavity center. The ellipse model considers the spatial variation of the stability limit, with its major axis aligning with the superior-inferior direction and its minor axis aligning with the posterior-anterior direction. The polynomial formulation also accounts for spatial variability. It was derived empirically from concavity compression tests [6] as a set of polynomial functions that fit eight discrete points along the glenoid border [14, 28].

A rapid muscle redundancy (RMR) solver was recently developed to solve the load sharing problem in the thoracoscapular model [31]. The RMR solver adopted a stability model that approximated the stability border as a circle, and estimated higher activation of the rotator cuff muscles than without any constraint, suggesting more reasonable active contribution of the rotator cuff muscles to GH stability. The GH-JCF vector, however, was frequently directed along the circular perimeter, as this was the “cheapest” solution. This finding raises questions to its validity in predicting muscle activity.

Instead of inequality constraint formulations that force the joint contact force to be directed within a specified area, a different strategy to model GH stability was presented by introducing a conditional penalty to the objective function that triggers if the GH-JCF direction crossed a specified stability perimeter [32]. The penalty term was proportional to the squared distance that the GH-JCF direction deviated outside the perimeter. This formulation, however, often estimated GH-JCFs that were directed along or occasionally outside the stability perimeter.

Several knowledge gaps exist with the current stability formulations. The existing inequality constraint models approximate the stability perimeter differently, and there is no clear evidence on which model is most reasonable. For most formulations, the directions of the estimated joint contact forces were evaluated and considered plausible if they fell within the perimeter, but were not compared to gold standard *in vivo* measurements. The exception to this is the ellipse model implemented into the Delft shoulder model, for which computed GH-JCFs were tested against *in vivo* measurements from the *Orthoload* dataset [17, 33]. The model reproduced the direction of the GH-JCF inside the glenoid cavity. Discrepancies were, however, present between the measured and estimated GH-JCF magnitudes, specifically for dynamic tasks above 90^*o*^ of shoulder elevation, which the authors attributed to the inability of the model to reproduce co-contraction [33]. GH-JCF magnitudes computed in the AnyBody implementation of the Delft shoulder model [17] were also compared to *in vivo* measurements, but not directions, which can be expected to be more sensitive to GH stability models [28, 29]. Implemented in a different shoulder model, the ellipse formulation was also shown to underestimate co-contraction of the rotator cuff muscles compared to *in vivo* fine-wire EMG measurements [8].

The objectives of this study were to determine how different formulations of GH stability in musculoskeletal simulation influence estimated GH-JCF magnitude and direction, in relation to reported *in vivo* joint contact force measurements, and to evaluate how rotator cuff muscles co-contraction was predicted with the different stability formulations.

## Materials and methods

We performed musculoskeletal simulations on the experimental data available from the *Orthoload* dataset. In the thoracoscapular model, joint angles, velocities and accelerations were computed via inverse kinematics. Muscle redundancy was then solved with the RMR solver repeatedly, for several different joint stability formulations that constrain the direction of the GH-JCF; these include constraining the stability perimeter as different shapes, then by penalizing joint contact force directions outside of a defined perimeter. Estimated joint contact force magnitude and direction were compared with *in vivo* measurements.

### Orthoload Dataset

Available *Orthoload* data from subject S8R with an instrumented hemiarthroplasty in the right shoulder (male, 74 y, mass 83 kg, height 171 cm) was used in this study [34–36]. The subject had previously undergone joint hemiarthroplasty with a delto-pectoral approach with no observed nerve or rotator cuff damage. The subject performed three tasks: lateral raise (abduction), posterior raise (backward flexion), and anterior raise (forward flexion) with a 2.4 kg handheld dumbbell. Kinematic data had been measured with marker clusters placed on thorax, scapula (on the acromion), upper arm, and forearm using Optotrak motion capture system (Northern Digital Inc., Canada) cameras at a sampling frequency of 50 Hz. Bony landmarks of the scapula: Angulus Acromialis, Trigonum Spinae, and Angulus Inferior were identified in calibration trials using a scapula locator [37], and constructed in dynamic trials from the acromion cluster. GH-JCFs were measured with the instrumented joint prothesis that sent data wirelessly to an external measuring system and resampled to 50 Hz. Forces were measured in the instrumented joint prosthesis coordinate frame of reference. A coordinate transformation from the prothesis frame to the local coordinate system of the glenoid cavity was then obtained from subject-specific computed tomography images of the participant.

### Musculoskeletal Model

The thoracoscapular model consists of a multibody chain that defines articulation among right upper body clavicle, scapula, humerus, ulna, and radius [25]. A closed chain mechanism is formed with a 4 degree-of-freedom (DoF) scapulothoracic *OpenSim* mobilizer [24], a 2 DoF joint between the thorax and clavicle, and a joint between the clavicle and the acromion that allows 3D rotation but no translation. The GH joint is modeled as a gimbal joint with 3 DoFs according to the ISB standard [38]. The elbow is modeled as a 1 DoF joint, and the forearm with a 1 DoF joint between the ulna and radius to model forearm pronation/supination. The hand is rigidly fixed to the radius and the whole multibody chain is anchored to the ground via a 6 DoF joint at the thorax. The model includes 33 muscle elements that represent core muscles of the shoulder girdle, including rotator cuffs, deltoids, serratus anterior, latissimus dorsi, among others. In this study, the model was used linearly scaled using the *OpenSim* geometric scaling tool then used to simulate all tasks. The subject’s handheld load was applied as a mass at the hand’s center of mass.

### Muscle Redundancy Solver

Described by Eq 1a, the RMR solver minimizes the sum of squared muscle activations at each time frame (static optimization), while adding a penalty on reserve actuators to ensure that motion is generated predominantly by muscles [31]. An equality constraint is imposed to match the measured 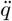 and estimated 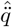 coordinate accelerations (Eq 1b), and the rate of change of muscle activation is restricted to respect the physiological first-order activation dynamics (Eq 1c).

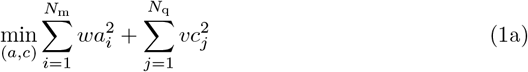

subject to

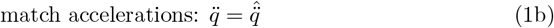

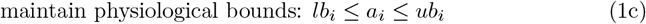

where *a*_*i*_ is activation of muscle *i, w* is the weight assigned to activation of muscle *i* which is chosen as one for all muscles, *c*_*j*_ control input to reserve actuator *j, v* is the weight assigned to control of reserve actuator *j* which is always equal to ten, *lb*_*i*_ and *ub*_*i*_ are the lower and upper physiological activation bounds, respectively, of muscle *i*, as described by Belli et al. [31].

### GH Stability Formulations

GH stability was formulated with different approaches, first as constraints that restrict the GH-JCF direction to lie within a specified perimeter, then instead by penalizing joint contact force directions in proportion to the deviation from a single point or from a defined perimeter (Fig 1). In all formulations, the direction of the joint contact force vector 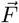 is described by the intersection point (*o*_*p*_ in Fig 2, and Fig S1 in S1 Appendix) of the force’s line of action with the glenoid plane—the plane containing the glenoid cavity border. This intersection point is referred to here as the GH-JCF direction on the glenoid plane, with coordinates described by the position vector 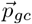, described by Eq S4 in S1 Appendix.

**Fig 1.**
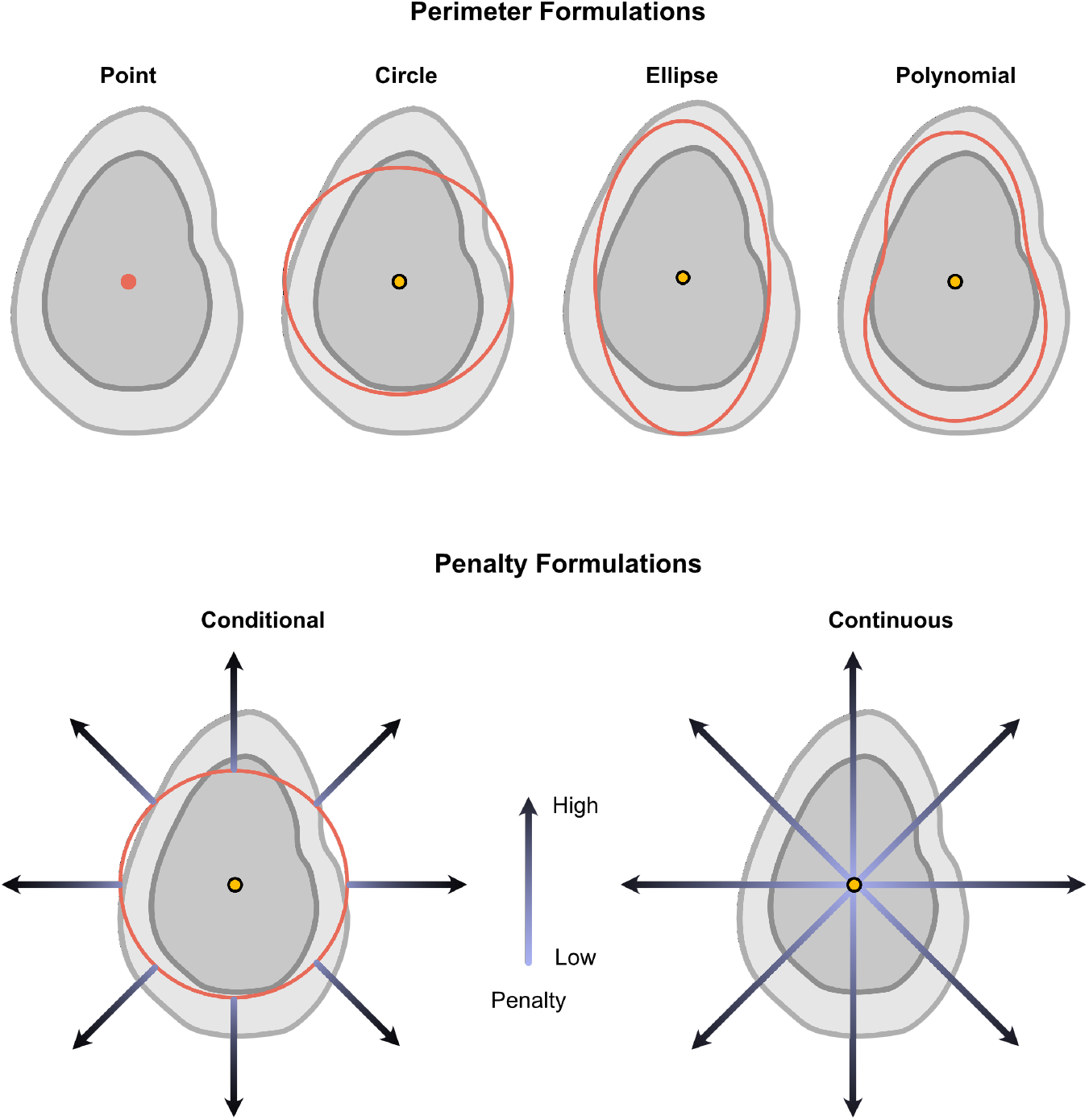
Demonstration of GH stability formulations. Image of glenoid cavity and the different perimeters for GH stability formulations investigated, not necessarily drawn to scale. The polynomial shape is fit through 8 discrete points reported from empirical concavity compression tests [6].

**Fig 2.**
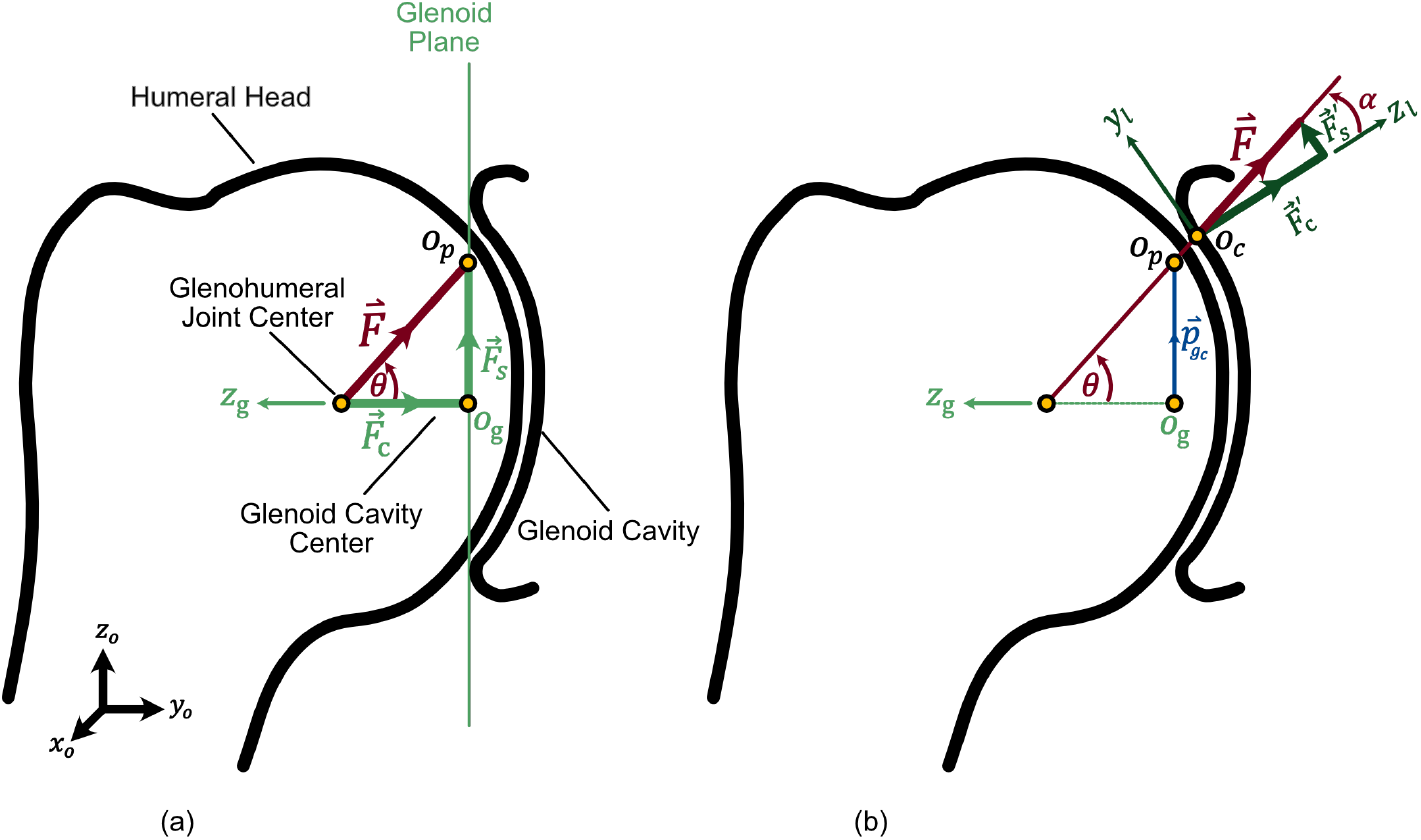
GH-JCF decompositions. Illustration of the GH-JCF vector 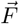. Two decompositions of 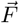 are demonstrated. (a) 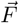 is decomposed onto the glenoid plane as a compressive component 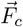 in the direction of the *z*_*g*_ *− axis* and a shear component 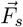 normal to the *z*_*g*_ *− axis*. (b) 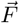 is decomposed as a compressive component 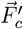 in the direction *z*_*l*_ normal to the curve, and a shear component 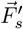along *y*_*l*_ in the direction tangent to the curve.

#### Point

In this formulation, the GH-JCF is forced to intersect the point at the glenoid cavity center. Described by Eq 2, this formulation forces the angle *θ* between 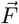 and the vector from the GH joint center to the glenoid cavity center (*z*_*g*_ axis) to be 0 (Fig 2).

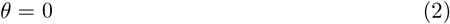

### Circle

In the RMR solver, GH stability is formulated as an inequality constraint on the GH-JCF direction on the glenoid plane, preventing it from lying outside a circular stability perimeter with a center at glenoid cavity center. We chose the diameter of this circle *D* such that the perimeter characterizes a stability limit of 0.5. A stability limit of 0.5 is chosen within the reported ranges of empirical stability limits (0.32 *−* 0.6) [7, 39].

The Circle formulation is described by Eq 3.

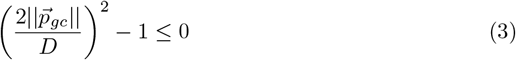

where, 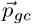 is a position vector that describes the direction of the GH-JCF on the glenoid plane (Fig S1, and Eq S4 in S1 Appendix). Note that, the stability limit is a dimensionless quantity however,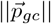, and *D* have the dimension of length.

#### Ellipse

In this formulation, the direction of the GH-JCF is constrained to stay within an elliptical perimeter, with a long axis in the superior-inferior direction and short axis in the anterior-posterior direction (Fig 1), as per previous studies [8, 10]. Eq 4 describes the inequality constraint that formulates the Ellipse model.

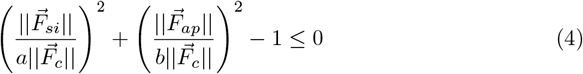

where *a* = 0.61 and *b* = 0.34 are the long and short axes of the ellipse that represent the empirical stability limits observed in concavity compression tests [6].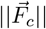 is the magnitude of the compressive component of the estimated GH-JCF that acts perpendicular to the glenoid plane (Fig 2, and Fig S1 in S1 Appendix). 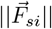 is the magnitude of the estimated shear force 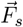 in the superior-inferior direction, and 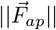 is the magnitude of 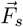 in the anterior-posterior direction. The shear force 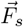 is the component of the estimated GH-JCF parallel to the glenoid plane.

#### Polynomial

In this formulation, the stability perimeter is defined by a 6th order polynomial fit of the empirical values reported from concavity compression experiments [6] (Fig 1). The formulation is implemented as an inequality constraint described by Eq 5.

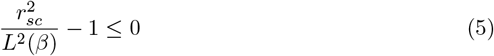

where *L*(*β*) is a 6^*th*^-order polynomial fit of the empirical stability limits reported by concavity compression tests (Fig 1) [6], and *β* is the radial angle in the glenoid plane from the center (Fig S1 in S1 Appendix). Described by Eq 6, planar shear-to-compressive ratio *r*_*sc*_ is the ratio between the shear and compressive components of 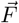 decomposed parallel and normal to the glenoid plane (Fig 2).

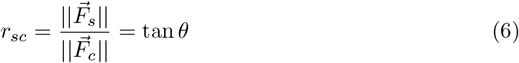

#### Conditional Penalty

In this formulation, a penalty is added as a cost in the objective function if the GH-JCF direction crosses the circular stability perimeter (Fig 1), again defined with radius that corresponds to a stability limit of 0.5. The penalty term is applied proportional to the planar shear-to-compressive ratio *r*_*sc*_ squared (Eq 6). The objective function in the original formulation Eq 1a is thus replaced by Eq 7.

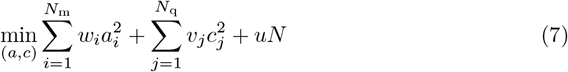

where *u* is the penalty weight, set a value of three, and *N* is a penalty term defined as follows,

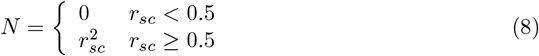

#### Continuous Planar Penalty

In this formulation, a penalty term is added to the objective function continuously, proportionally penalizing the distance squared from the GH-JCF direction away from the center of the glenoid cavity. The penalty term is equal to the planar shear-to-compressive ratio (Eq 6), and scaled by a weighting factor of *u* = 3. With the Planar Penalty formulation Eq 9 replaces the original objective function in the RMR solver Eq 1a.

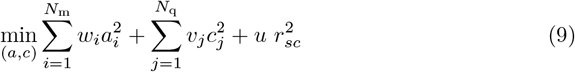

#### Continuous Curve Penalty

This formulation is similar to the Continuous Planar Penalty, but considers the shape of the glenoid cavity curvature by applying a penalty term that is equal to the curve shear-to-compressive ratio 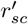 (Eq 10). This ratio is defined as the magnitude ratio between the components of 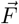 tangent and normal to the glenoid cavity curvature (Fig 2).

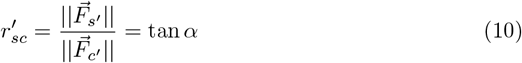

The curvature of the glenoid cavity was assumed to be circular with a radius and a center location that achieve specific height (4 *cm*) and depth (2 *mm*) of the glenoid cavity (Fig S2 in S1 Appendix) based on *in vitro* morphological measurements [44]. With this formulation, the original objective function Eq 1a is replaced by Eq 11. The penalty term in Eq 11 scaled by a weighting factor of *u* = 3.

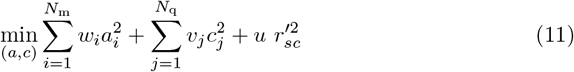

#### None

Muscle redundancy was also solved with no inequality constraints or cost penalties stability formulations, referred to as the None model.

### Simulation Pipeline

Inverse musculoskeletal simulations were performed on the *Orthoload* dataset using the unscaled thoracoscapular model. Muscle redundancy was solved for all tasks with the RMR solver and the relevant GH stability formulation..

### Sensitivity Analysis

Sensitivity analysis was performed to evaluate the effects of the stability limit value on the solution obtained from the inequality constraints formulations. A case study was performed on the Polynomial model by scaling up and down the size of the perimeter by a factor of 2, with results shown in S1 Appendix. Sensitivity analysis was also performed on the weights of the penalties in the objective function in the penalty formulations; weights were assigned *u* = 1, *u* = 3, and *u* = 10 in the RMR solver.

## Results

### GH-JCF Magnitude

With no stability formulation (None model), the overall pattern of the GH-JCF magnitude was similar to the measured force in all tasks (Fig 3). Predicted peaks were, however, occasionally temporally shifted, particularly in lateral and anterior raises. In lateral and anterior raises, the None model underestimated the magnitude over the majority task duration, most notably in lateral raise. In posterior raise, however, the None model reproduced high magnitudes compared to the *in vivo* measurements most of the time.

**Fig 3.**
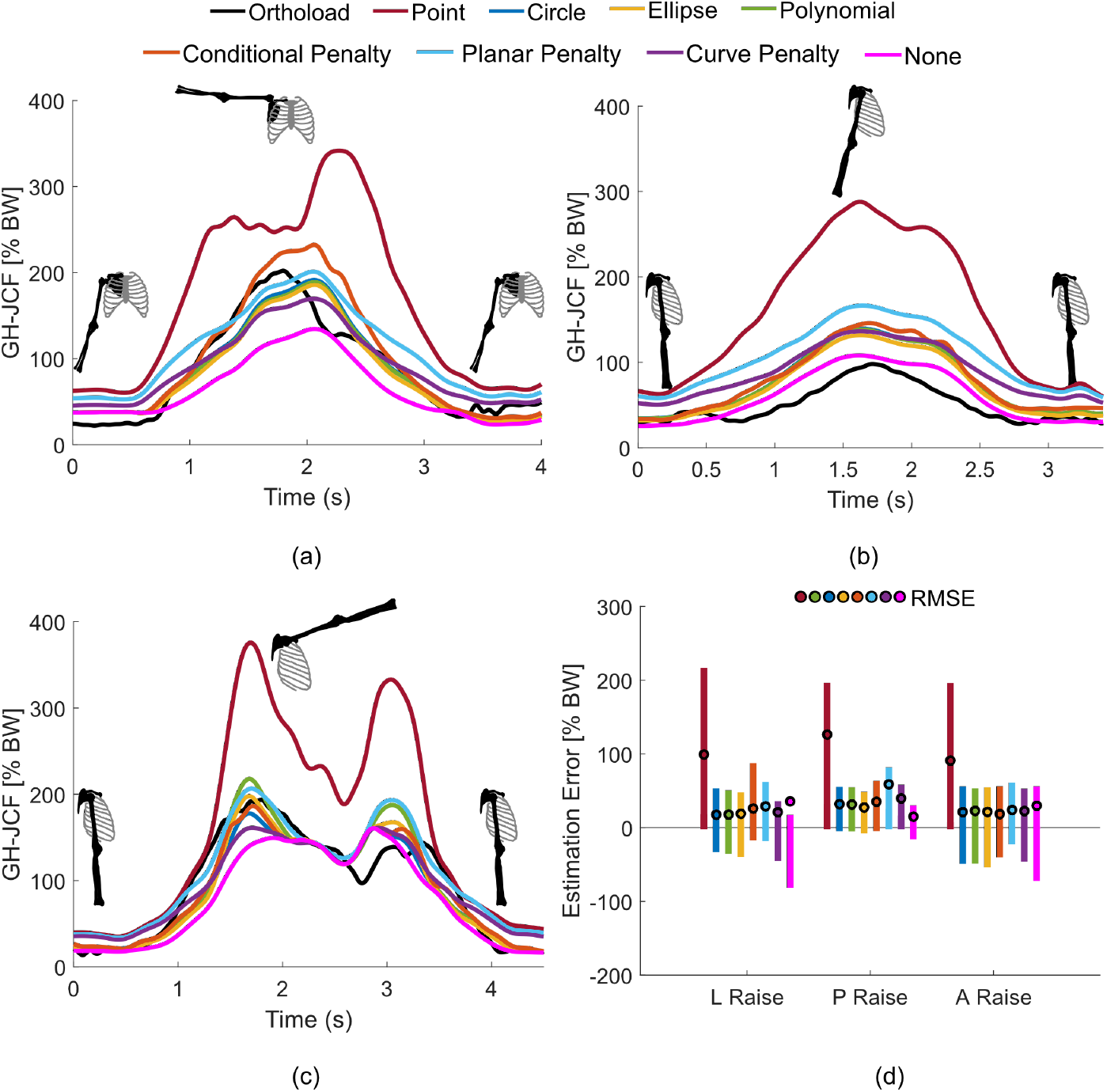
Estimated GH-JCF magnitude vs *in vivo* measurements. (a) - (c) GH-JCF vector magnitude as a percentage of the body weight (%BW) in all tasks with the different stability formulations. (d) Estimation error in GH-JCF magnitude compared to *in vivo* measurements. Bars show the maximum positive (overestimation) and minimum negative (underestimation) errors. Circles indicate the root mean square error (RMSE) between the measured and estimated magnitudes throughout each task. L Raise (Lateral Raise) P Raise (Posterior Raise), A Raise (Anterior Raise).

All perimeter formulations predicted GH-JCF magnitudes that agreed reasonably well with the *in vivo* measurements, although the timing of the peaks was occasionally shifted. With these formulations, prediction of the magnitude was closer to the measurements than the None model in lateral and anterior raises. In posterior raise, however, the perimeter models overestimated the magnitude more than the None model. In all tasks, the Point model overestimated the GH-JCF magnitude the most, with approximately 105% body weight (BW) root-mean-square-error (RMSE) compared to the measured force magnitudes. Other perimeter models estimated force magnitudes with RMSE approximately around 18% BW in lateral raise, 30% BW in posterior raise, and 22% BW in anterior raise.

The penalty formulations predicted GH-JCF profiles similar to the perimeter formulations in all tasks, with somewhat higher RMSEs, most notably in posterior raise with the Planar Penalty (59% BW). In lateral and anterior raise, RMSEs with the penalty formulations were approximately 25% BW, and 21% BW, respectively.

Among all stability models, predictions with the lowest RMSEs were computed with the Circle model in lateral raise (*≈* 18% BW), the None model in posterior raise (*≈* 15% BW), and the Conditional Penalty model in anterior raise (*≈* 19% BW).

### GH-JCF Direction

During all tasks, the *in vivo* measured GH-JCFs were always directed inside the glenoid cavity, often slightly posteriorly, though not always (Fig 4). Except for some time frames at the beginning and end of lateral and anterior raise tasks, with the None model, the GH-JCF direction was directed outside of the glenoid cavity—most notably up to 13 *cm* superior and 7 *cm* anterior to the glenoid in the lateral raise. With the Point formulation, the estimated GH-JCF was directed precisely at the glenoid cavity center. With the Circle, Ellipse and Polynomial formulations, the GH-JCF was directed near or directly along the superior perimeters throughout all tasks.

**Fig 4.**
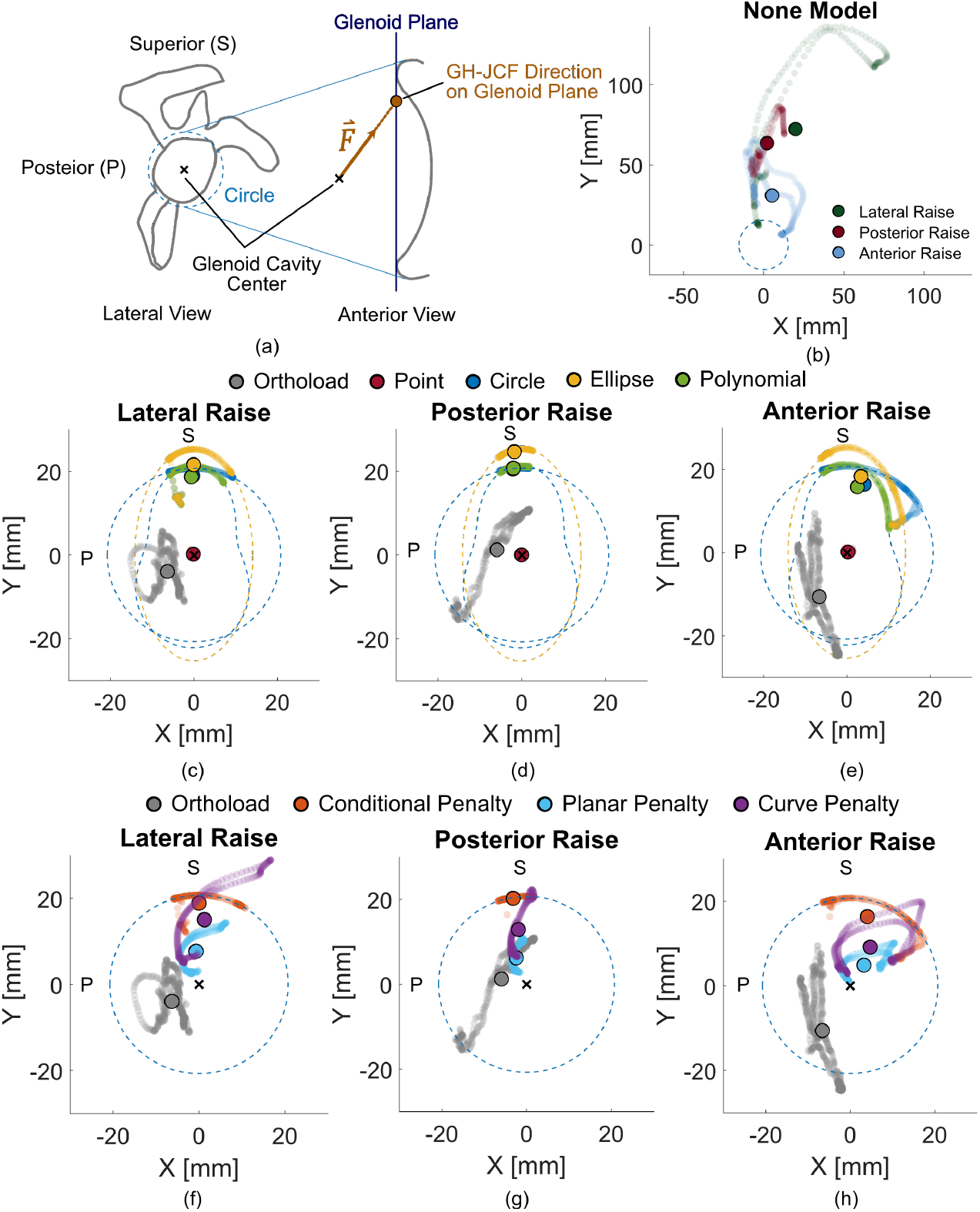
Estimated GH-JCF direction vs *in vivo* measurements. Direction of the GH-JCF vector on the glenoid plane during all tasks, measured *in vivo* and estimated with the different stability formulations. (a) Lateral and anterior views of the scapula, with the Circle stability perimeter illustrated for scale, (b) GH-JCF direction on the glenoid plane with no stability formulation (None). (c)-(e) GH-JCF direction on glenoid plane from different perimeter formulations. (f)-(h) GH-JCF direction on glenoid plane with the different penalty formulations. Circles show the mean projection during each task. Dotted lines represent the different stability perimeters of each stability formulation.

With the Conditional Penalty formulation, the GH-JCF direction was again often directed close to or along the superior perimeter. With the Continuous Penalty formulations, the GH-JCF direction intersected the glenoid more centrally in the glenoid, and agreed best with the *in vivo* measurements, wherein estimations with the Planar Penalty matched slightly better than with the Curve Penalty.

### Rotator Cuff Muscle Activation

Estimated rotator cuff muscle activations were notably higher with any stability formulation than in the None model (Fig 5), with highest activations from the Point model. In all tasks, activation was higher with the penalty models than with any of the shape perimeter models. Among the penalty models, rotator cuff activations were higher in all tasks with the Planar model.

**Fig 5.**
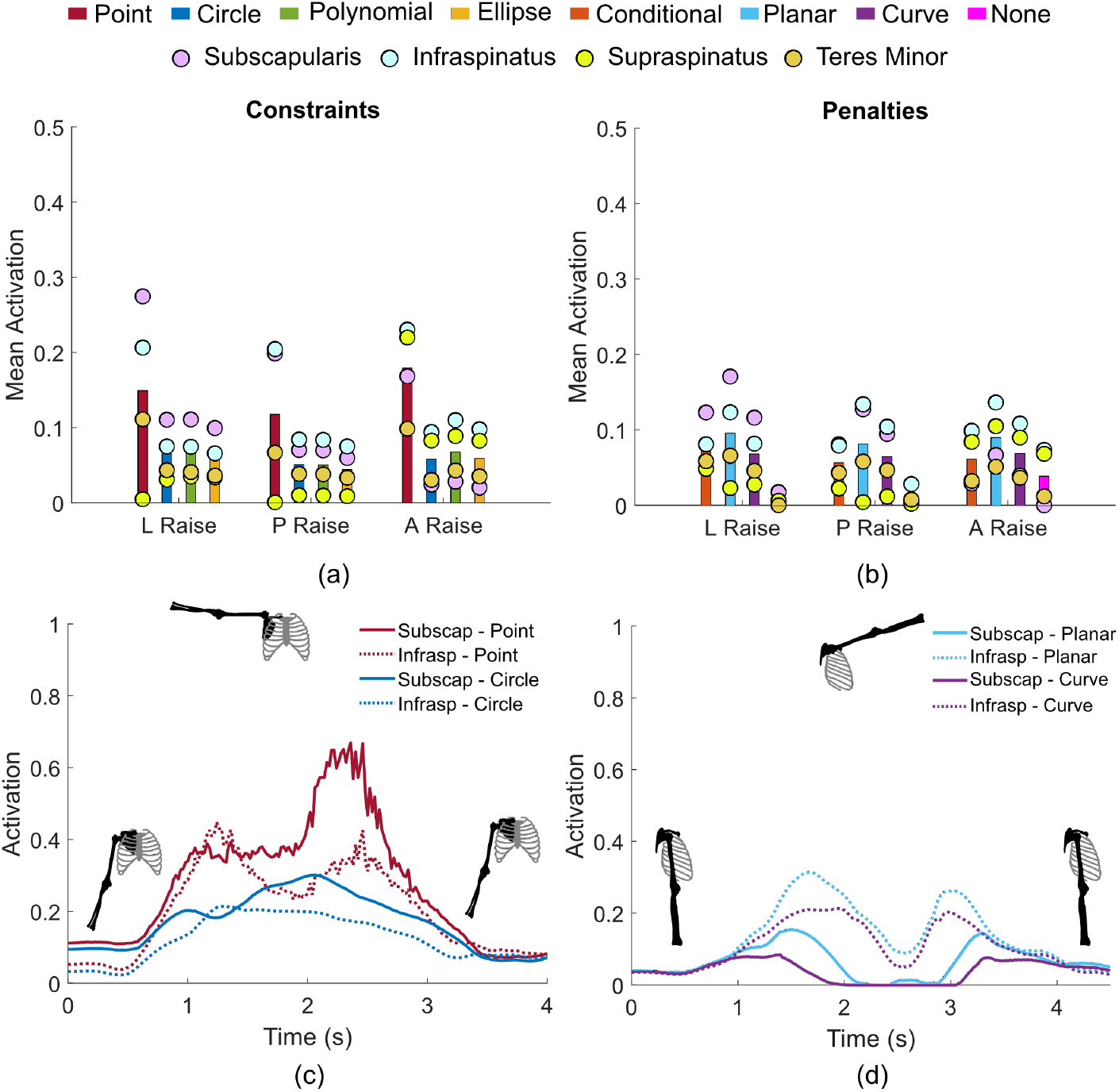
Estimated rotator cuff activation. (a), (b) Mean activation of the rotator cuff muscles group in all tasks with the perimeter and penalties formulations. Circles show the mean activation of each individual muscle throughout each task. Bars show the mean activation of all muscles. (c), (d) Activation of selected rotator cuff muscles with different stability formulations in lateral and anterior raise tasks. Subscap (Subscapularis), Infrasp (infraspinatus).

Activation levels of the rotator cuff muscles varied between tasks. In the lateral raise, the subscapularis muscle had the highest activation with all stability models, followed by the infraspinatus and the teres minor. In the posterior raise, the infraspinatus muscle had the highest activation, followed by the subscapularis and the teres minor with all stability model except the Conditional Penalty model where the subscapularis came first. In the anterior raise, the infraspinatus muscle had the highest activation, followed by the supraspinatus and the teres minor with most of the stability models. Activation patterns also varied across stability formulations. In the anterior raise, the teres minor muscle was the least activated muscle with the Point, the Curve Penalty and the Planar Penalty models, and was the third most activated with all other models.

### Computational Efficiency

Computational efficiency on an Acer Predator Triton 500 SE laptop with Core i7-11800H processor varied with stability models (Fig S3 in S1 Appendix). In lateral raise for example, computational time ranged from 43.2 seconds for the None model to 69.1 seconds for the Point model. In the posterior raise, times ranged from 39.2 seconds to 54.8 seconds. Overall, differences in computational time between perimeter and penalty formulations remained within 15 seconds.

### Sensitivity Analysis

We evaluated the sensitivity of the solutions reproduced by the different formulations to the size of the stability perimeter and the value of the optimization weight *u* (Fig 6, and Fig S4 in S1 Appendix). Results showed that reducing the diameter of the stability perimeter resulted in stricter predictions with an increased GH-JCF magnitude and closer direction to the glenoid cavity center, whereas increasing the diameter of the stability perimeter had the opposite effect. Likewise, increasing the optimization weight led to a higher GH-JCF magnitude and closer direction to the glenoid cavity center, while decreasing it resulted in a lower GH-JCF magnitude with a direction farther from the glenoid cavity center. It was simultaneously observed that, by reducing the size of the stability border and increasing the optimization weight, the activation of the rotator cuff muscles increased, and vice versa.

**Fig 6.**
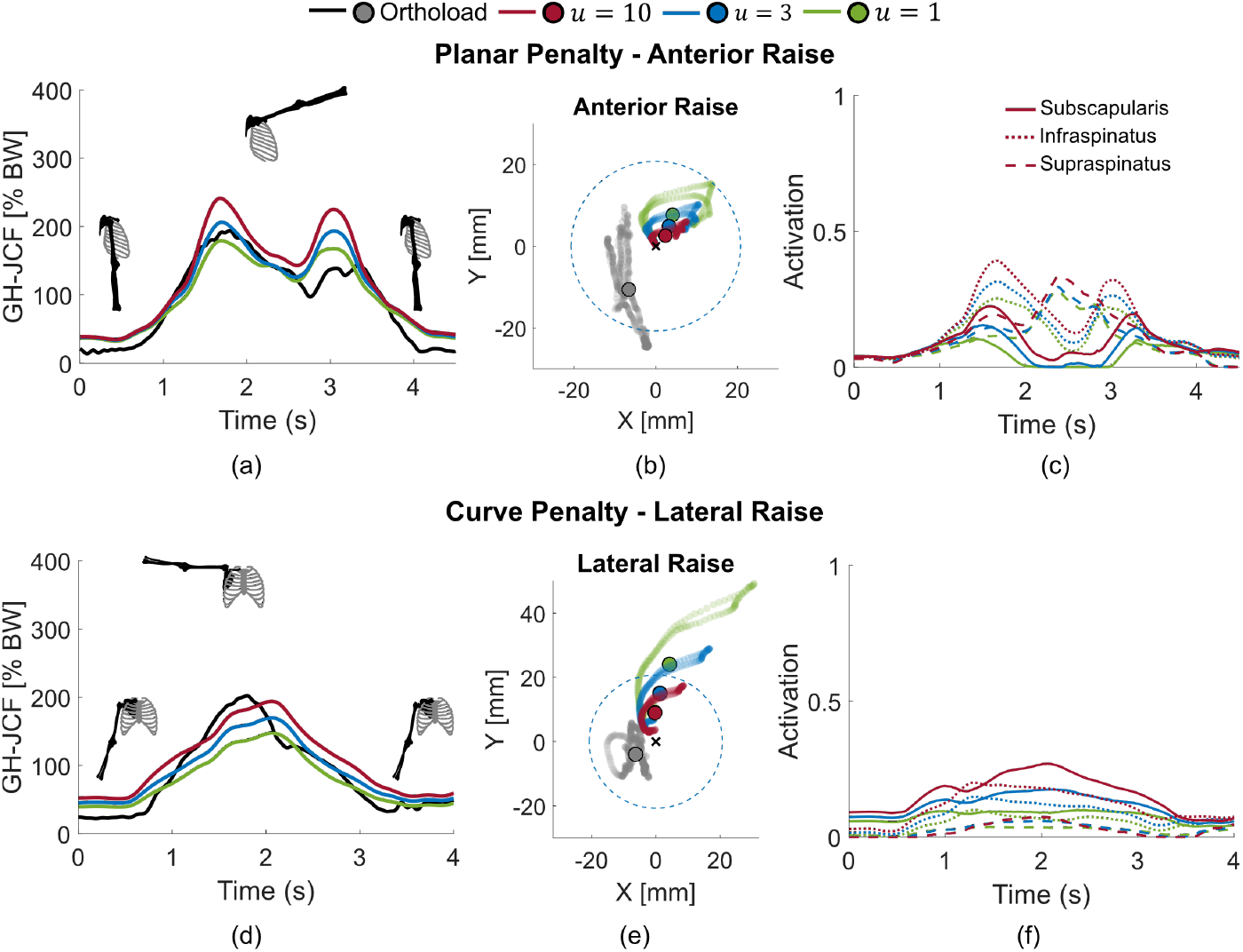
Sensitivity analysis. Estimated joint contact forces, GH-JCF direction on the glenoid plane, and activation of the rotator cuff muscles in anterior raise task with the Planar Penalty formulation (top row), and in lateral raise task with the Curve Penalty formulation (bottom row), for varying penalty weights *u* in the cost function. (a), and (d) show the GH-JCF magnitude. (b), and (e) GH-JCF direction on the glenoid plane. Circles show the mean of the GH-JCF direction on the glenoid plane over time; light shades show the trajectory of the GH-JCF direction with time; dotted lines represent the Circle stability border. (c), and (f) Activation of the rotator cuff muscles.

## Discussion

In this study, we investigated different approaches to model GH stability in musculoskeletal simulations using the thoracoscapular shoulder model and the RMR solver. The thoracoscapular *OpenSim* model incorporates a validated model of the scapulothroacic articulation. The RMR solver includes a formulation of the GH stability that can be customized. Both are freely available. The goal of this investigation was to determine which GH stability formulations most faithfully reproduced GH-JCF magnitude and direction measured with an instrumented shoulder prosthesis.

Existing formulations of GH stability have either shown limited accuracy or lack quantitative validation. We quantitatively investigated and tested different GH stability formulations via two general schemes—incorporating inequality constraints that constrain the GH-JCF direction to different perimeters, and introducing penalties in the objective function that allow but penalize GH-JCF directions that deviate from the glenoid cavity center. Estimated GH-JCFs without any GH stability formulation (None) were lowest in magnitudes and directed several centimeters above the glenoid cavity during the three tasks, which in reality would likely lead to a dislocated shoulder. With the strictest stability formulation (Point), that forced the estimated GH-JCF to be directly in the center of the glenoid cavity, GH-JCF magnitudes were overestimated by up to 2 times body weight during the movements, achieved by excessive muscle activations and co-contractions. This formulation can thus also be considered inaccurate.

Constraining the GH-JCF direction within specified perimeters using inequality constraints estimated GH-JCF that, though closer to the glenoid than with no constraints, still did not agree with those measured experimentally; they were instead nearly always directed along the superior perimeter, and can likewise be considered unrealistic. The Conditional Penalty formulation was constructed as a sort of compromise, as it allowed GH-JCF to be directed outside of the defined perimeter, but at an additional cost in the objective function. However, largely the same behavior was found; GH-JCF direction was still directed very close to or along the superior perimeter throughout most of the activities, so it can also be considered inaccurate.

The Continuous Penalty formulations have no specified perimeter or border, but instead encourage the GH-JCF to be directed centrally in the glenoid cavity, achieved by penalizing the GH-JCF direction that deviates from the glenoid cavity center, equally in all directions. Both continuous penalty formulations estimated GH-JCF magnitudes and directions that agree rather well to those measured *in vivo*. They also encouraged enough muscle co-contraction of particularly the rotator cuff muscles to achieve GH stability, though no fine wire EMG data was available in the Orthoload data set for a direct comparison. Taking into account the curvature of the glenoid cavity (Curve penalty) vs. simplifying the glenoid to a plane (Planar penalty) did not result in substantially more accurate GH-JCF magnitudes or directions, suggesting that the more simplified geometric model in the muscle redudancy solver may suffice.

Interestingly, in the posterior raise, the estimated GH-JCF magnitude was overestimated in all stability formulations, most so in the strictest formulation and least in the None model. This overestimation is likely attributable to the absence of ligament forces in the current model. In posterior raise movements, several ligaments in the shoulder joints likely contribute to GH stability, specifically the coracohumeral ligament between the the humeral head and the coracoid process of the scapula and the superior GH ligament between the humeral head and the glenoid cavity [1]. If ligaments were included in the model, it can be expected that RMR solver would predict lower muscle activations and co-contractions, as ligament force is passive and would be ”cheaper” metabolically.

Estimated GH-JCF directions were most accurate with the two Continuous Penalty formulations. They do not, however match *in vivo* measurements exactly, nor can they be expected to, for several reasons. First, the modeled GH joint is simplified; it ignores humeral head translation and the effect of the glenoid labrum on stabilizing the GH joint. *In vitro* studies showed that GH stability is spatially dependent on the point of contact between the humeral head and the glenoid cavity as well as on the thickness of the glenoid labrum at that point [6, 7]. Inverse simulation using skin markers measurements is furthermore not likely suitable to accurately reproduce humeral head translation, which is on the same order of magnitude of measurement errors arising from soft tissue artifacts, particularly in scapular movement [40, 41]. Second, while simulations were performed with a geometrically scaled model, additional scaling based on muscle strength might have improved model predictions. This, however, was not possible in the current study, as no information about the subject’s strength was available in the dataset, nor was fine wire EMG data. Third, the subject underwent a shoulder hemiarthroplasty, so his joint topology and musculoskeletal system will not precisely match even a scaled model.

With the Curve Penalty model, the predicted direction of the GH-JCF on the glenoid plane was slightly further from the glenoid cavity center than with the Planar Penalty model. This is because the shear force magnitude decomposed tangent to the glenoid curve is always less than or equal to the shear force magnitude projected onto the glenoid plane. This means that the penalty term in the Curve Penalty formulation will be smaller than in the Planar Penalty formulation for any non-zero angle *θ* and similar weight *u*, and the Curve Penalty formulation will predict less co-contraction. It is, however, worth mentioning that the two models can equivalently lead to the same behavior by tuning the weight of the penalty term *u*, as was indicated in the sensitivity analysis.

The GH-JCFs have been estimated via inverse musculoskeletal simulation in a few published studies, one using the OpenSim version of the Delft shoulder model [42] and another other using the Newcastle shoulder model, which defined muscles similarly to the Delft model [16]. In both of these studies, the estimated GH-JCFs were directed within the glenoid cavity, even without applying any shoulder stability models. These findings contradict previous findings by van der Helm who, using the Delft shoulder model, reported that a stability perimeter was necessary in tasks like lateral and anterior raise [43]. It is worth mentioning that the musculoskeletal models used in these studies characterized joints and muscles slightly differently than the thoracoscapular model in the current study. Future work can investigate this issue by testing the proposed stability formulations on the OpenSim version of the Delft and/or Newcastle shoulder models.

Some other limitations exist with this study. The participant underwent a surgery that sacrificed the joint capsule, so it is likely that greater than normal muscle forces were required to compensate for the absence of the capsule’s stabilizing effect. This limitation is unavoidable in this gold standard method validation, as it requires surgical implantation of an instrumented joint prosthesis. Data measured *in vivo* that includes both forces and fine-wire EMG of deep muscles would be of some use in further validation studies, but to the best of our knowledge, no such data exists.

The subject had a prosthetic humeral head but an intact glenoid, likely altering the GH joint geometry and muscle moment arms. These geometric changes were not accounted for and may have influenced the predictions, though likely to a minor extent. For the perimeter formulations, the GH-JCF direction is always constrained to the defined perimeter, as confirmed by the sensitivity analysis; even when the perimeter diameter was larger or smaller, the GH-JCF direction remained on the perimeter. Similarly, in Continuous Penalty formulations, the formulation will still penalize deviations of the GH-JCF direction proportionally from the glenoid cavity center. Even with more personalized models of muscle moment arms, discrepancies between measured and predicted values would likely exist due to model simplifications, such as the omission of humeral head translation. While GH-JCF magnitudes might also depend on geometric variations, the overall pattern and trends would likely remain consistent, closely approximating measured values.

It should also be noted that some gap filling of marker trajectories in the Orthoload dataset was required. These gaps were interpolated using a spline fit, which sometimes introduced artifacts, particularly in trunk markers at the beginning or end of movements. To minimize these effects, affected markers were either disregarded when possible or assigned lower weights in the inverse kinematics solution.

Finally, data from only one subject was used for model validation; data from a greater number of subjects would have strengthened our model formulation, but likely not by much, as the predicted GH-JCFs still do not identically match the observed ones.

Future model improvements should include adding ligaments to the model, and improving muscle geometry models of the biceps and triceps, particularly for activities that involve elbow flexion. In the current model, biceps and triceps muscles do not have a wrapping point/surface around the elbow joint and are thus only suited for tasks with an extended elbow.

## Conclusion

This work comprehensively explored different formulations of GH stability in inverse musculoskeletal simulations using the thoracoscapular shoulder model and the rapid muscle redundancy solver. Estimates of GH-JCF direction and magnitude with the different stability formulations were tested against *in vivo* measurements. All stability formulations predicted the GH-JCF magnitude with relatively comparable accuracy, except for the strictest Point formulation, which constrained the joint contact force direction to the center of the glenoid cavity and overestimated the JCF magnitude. Penalty formulations that penalized deviations of the GH-JCF direction from the glenoid cavity center predicted GH-JCF direction more accurately than the other models, estimating GH-JCF within, rather than along or outside of, the glenoid cavity perimeter. These findings support the proposed stability formulations as benchmark methods for analyzing shoulder loading, providing a more adequate and reasonable description of GH stability.

## Supporting information

**S1 Appendix. Supplementary Material**. This appendix includes derivation of the GH-JCF direction on the glenoid plane and GH-JCF decompositions. It also includes supporting figures demonstrating sensitivity analysis, geometry of the glenoid cavity curvature adopted in the Curve Penalty formulation, and reserve actuator moments compared to joint moments.

## Acknowledgments

This work was funded by the Promobilia Foundation (ref. 18200, ref. 23300, and ref. A23165), and the Swedish Research Council (ref. 2018-00750), Stockholm, Sweden.

## Supplementary Material

### S1 Glenohumeral Joint Contact Force Direction on the Glenoid Plane

The glenohumeral joint contact force (GH-JCF) is a sliding vector (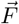 in Fig S1) that passes through the center of the glenohumeral (GH) joint, coincident with the center of the humeral head at any instant in time. The line of action of 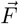 intersects with the glenoid cavity at a point. This point characterizes the GH-JCF direction in the glenoid cavity. The direction of 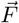 is used as a metric to assess the GH stability. The GH joint is stable as long as the direction of 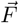 is within the glenoid border. Ideally, the intersection point of 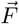 with the glenoid cavity curvature (point *o*_*c*_ in Fig S1) should lie on the articular surface of the glenoid. We, however, approximate this point by taking the intersection of line of action with the glenoid plane (point *o*_*p*_ in Fig S1), the *x*_*g*_*y*_*g*_ plane. The glenoid plane is the plane containing the border of the glenoid cavity. Axes *x*_*g*_ and *y*_*g*_ are the axes of the glenoid cavity reference frame which is defined as follows,

- *o*_*g*_: at the glenoid cavity center,
- *z*_*g*_: from glenoid cavity center to the GH joint center,
- *x*_*g*_: normal to *z*_*g*_ and lies in the plane containing the the humeral head center, the glenoid cavity center and the glenoid cavity edge, i.e., the glenoid plane,
- *y*_*g*_: normal to *x*_*g*_ and *z*_*g*_.

Bony landmarks were determined by visually locating them in the mesh geometry of model. Then, *OpenSim* model markers were placed at the identified landmarks to retrieve their position throughout the analysis. The coordinates of the intersection point *o*_*p*_ on the glenoid plane (i.e., GH-JCF direction on the glenoid plane) can be described by a vector 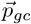, where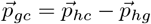. Vector 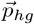 is the position vector from the GH joint center to the glenoid cavity center, and 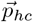 is the position vector from the GH joint center to the intersection point *o*_*p*_. The position vector 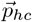 can be found as,

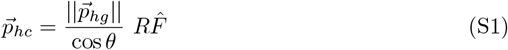

where, 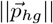 is the distance from the GH joint center to the glenoid cavity center, 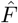 is a unit vector along 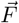 found as 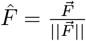, and *θ* is the angle between 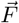 and 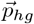 found as follows,

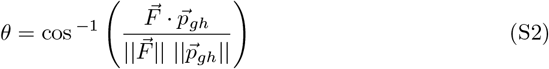

The matrix *R* is the rotation matrix that transforms from the global frame *o − x*_*o*_*y*_*o*_*z*_*o*_ to the glenoid frame *o*_*g*_ *− x*_*g*_*y*_*g*_*z*_*g*_. The rotation matrix *R* is defined as,

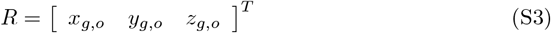

where *x*_*g,o*_, *y*_*g,o*_, and *z*_*g,o*_ are the unit vectors of the glenoid frame *o*_*g*_ *− x*_*g*_*y*_*g*_*z*_*g*_ defined in the global frame *o − x*_*o*_*y*_*o*_*z*_*o*_. The direction of 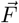 on the glenoid plane expressed in *o*_*g*_ *− x*_*g*_*y*_*g*_*z*_*g*_ is, thus, described by the position vector 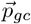 as follows,

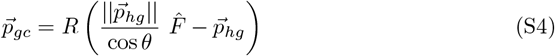

### S2 GH-JCF Decompositions

Two decompositions of the GH-JCF namely the planar decomposition and the curve decomposition are introduced. The planar decomposition decomposes 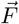 into shear and compressive components of in the parallel and normal directions to the glenoid plane (*x*_*g*_*y*_*g*_ plane) respectively (Fig S1). The shear force is the component of 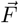 in the glenoid plane *x*_*g*_*y*_*g*_ plane 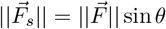 sin *θ*, and the compressive force is the component of 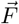 perpendicular to the *x*_*g*_*y*_*g*_ plane 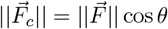 cos *θ* (Fig S1).

The curve decomposition decomposes 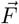 two components 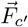, and 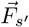 in the normal *z*_*l*_ and the tangent *y*_*l*_ directions to the curvature of the glenoid cavity at point *o*_*c*_ (Fig S1). Point *o*_*c*_ is the intersection of the line of action of 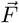 with the curvature of the glenoid cavity. We approximate the curvature of the glenoid cavity as a sphere with a radius 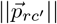 and a curvature center located on the *z*_*g*_ axis at a distance 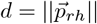 from the glenohumeral joint center. The radius of the curvature was chosen to achieve a specific height and width of the glenoid cavity (Fig S2) based on *in vitro* morphological measurements [1].

The normal component of 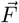 to the curvature is defined as 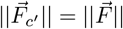 cos *α*, and the tangent component to the curvature is defined as 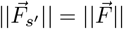 sin *α*. Angle *α* is the angle between 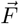 and 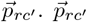 is the position vector from the curvature center to point *o*_*c*_. The magnitude of 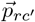 is equal to the radius of the curvature of the glenoid cavity. To find angle *α*, we apply the cosine law on angle *θ*^*′*^ which is the supplementary angle of *θ*. Applying the cosine law on *θ*^*′*^ results in a quadratic equation whose positive root is the length 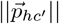 as described in Eq S5.

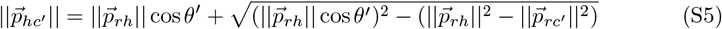

where 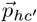 is the vector from the humeral head center to point *o*_*c*_, and 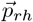 is the vector from the curvature center to the humeral head center. Angle *θ* is an exterior angle to the triangle whose sides are 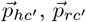, and 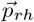. Therefore, angle *α* can be computed as follows,

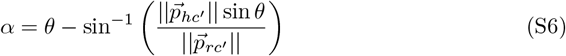

---

**Fig S1.**
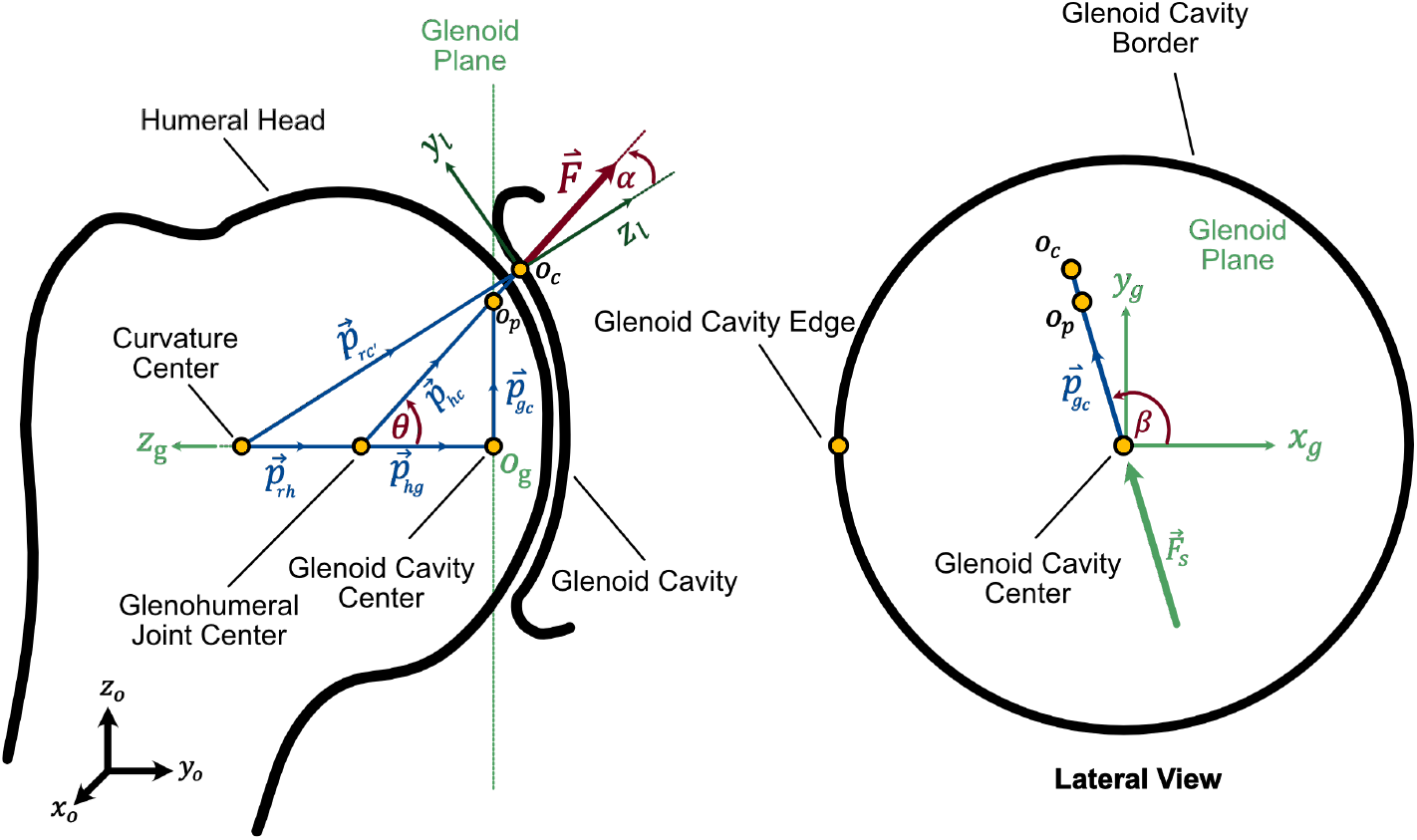
Demonstration of the direction of the GH-JCF vector on the glenoid plane. Point *o*_*p*_ is the intersection of the line of action of 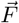 with the glenoid *x*_*g*_*y*_*g*_ plane. Axes *x*_*g*_ and *y*_*g*_ are the axes of the glenoid frame *o*_*g*_ *− x*_*g*_*y*_*g*_*z*_*g*_ whose origin is at the glenoid cavity center (GCC). Frame *o* − *x*_*o*_*y*_*o*_*z*_*o*_ is the global reference frame in which vectors are initially described.

**Fig S2.**
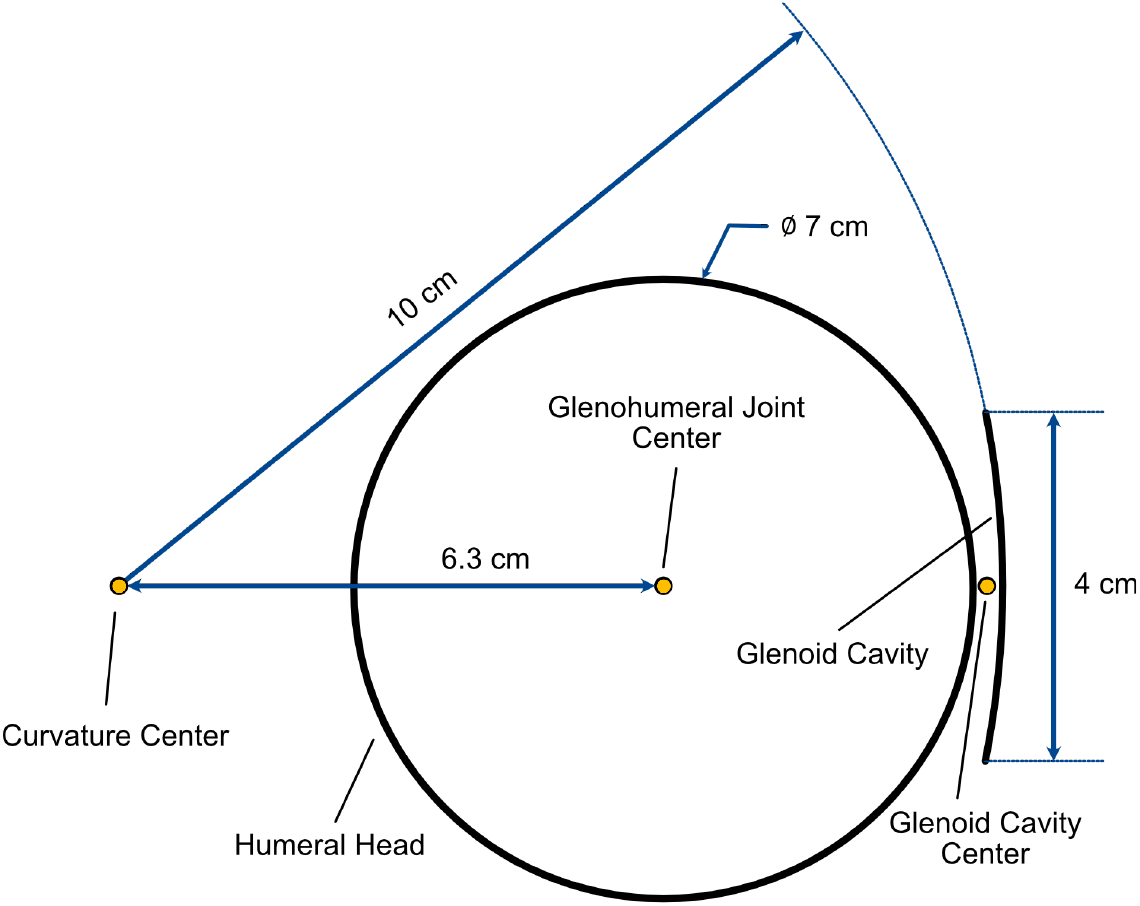
Geometry of the glenoid cavity curvature used in the Curve Penalty formulation. The glenoid cavity center is first found by placing a model marker roughly in the same plane of the glenoid cavity border on the mesh geometry. The height of the glenoid cavity is set equal to 4*cm* based on morphological measurements from *in vitro* experiments [1]. Then a circle is constructed such that its center is collinear with the glenohumeral joint center, and its radius achieve a glenoid cavity depth of 2*mm*—a value within the reported ranges from *in vitro* measurements [1]. The radius of the humeral head is constructed based on measurements from the model’s mesh geometry.

**Fig S3.**
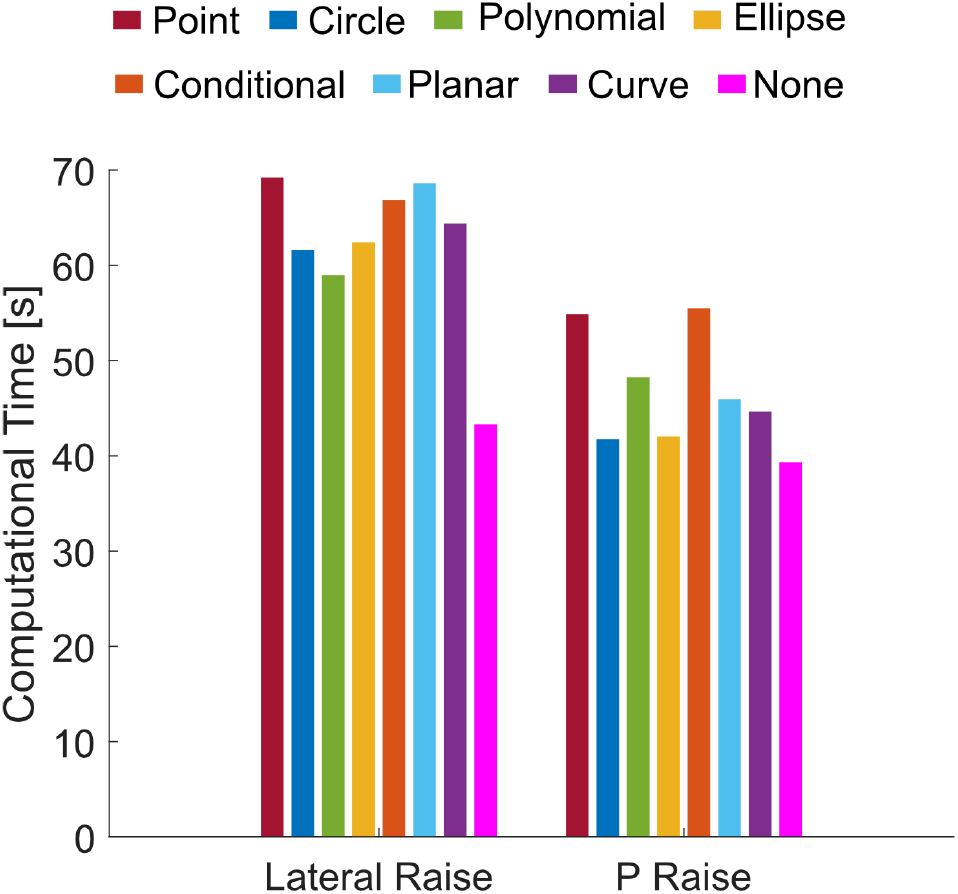
Computational time taken by the RMR solver to solve the entire task for each stability formulation in lateral and anterior raise tasks.

**Fig S4.**
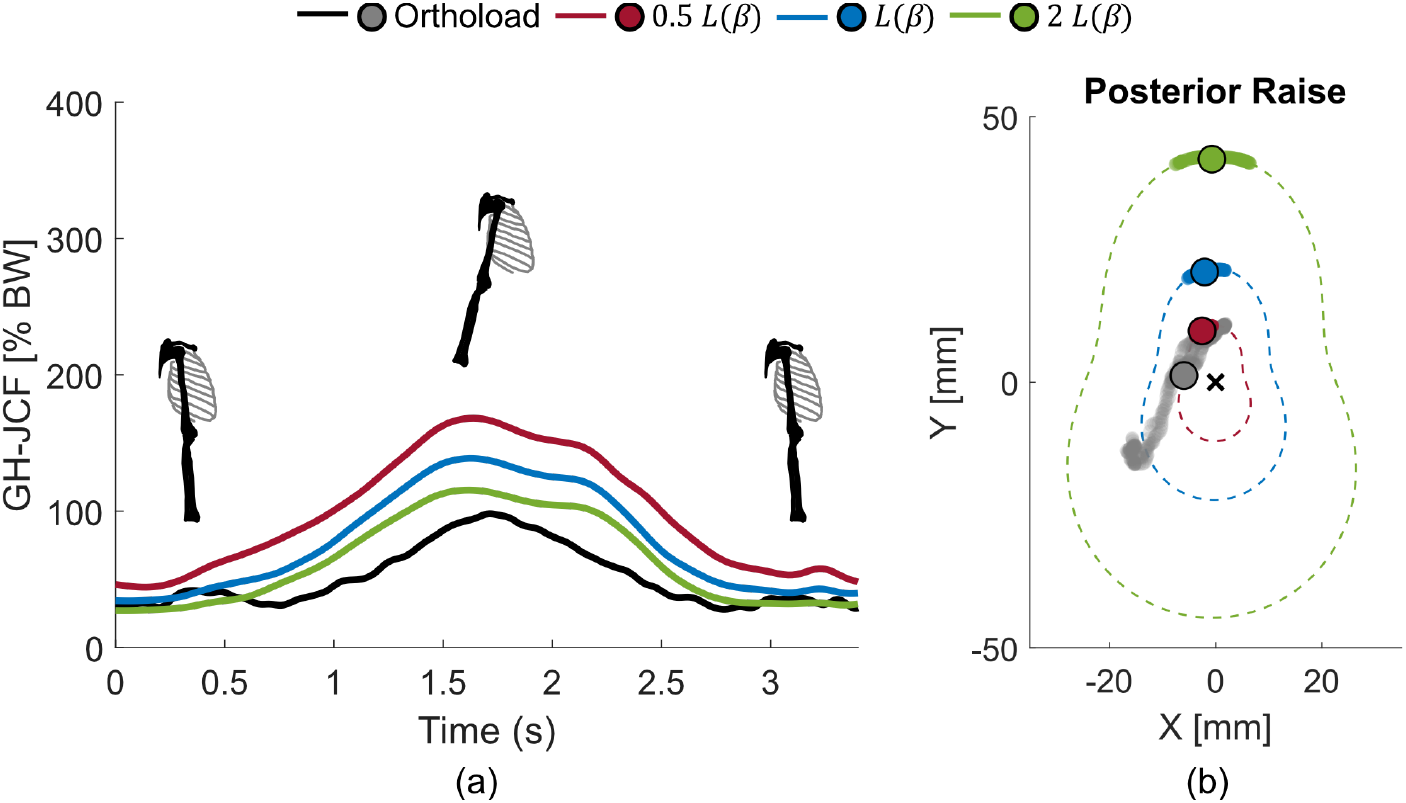
Estimated GH-JCF magnitude (a), and direction on the glenoid plane (b) in posterior raise task with the Polynomial formulation, for varying sizes of the stability perimeter. Circles show the mean of the GH-JCF direction on the glenoid plane over time; light shades show the trajectory of the GH-JCF direction with time; dotted lines represent the Polynomial stability border.

**Fig S5.**
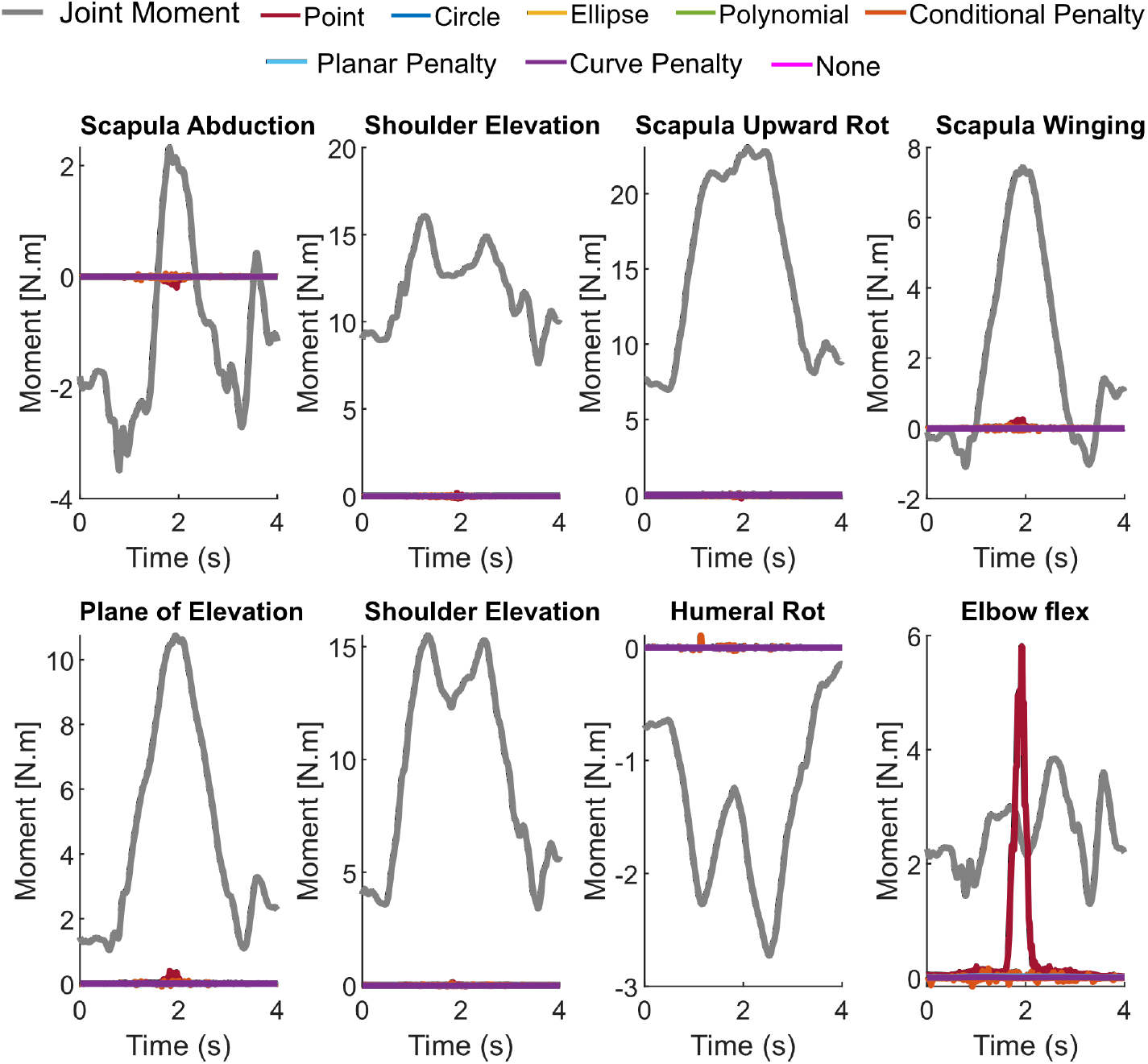
Moments of the scapulothoracic joint and the GH joint compared to moments of reserve actuators with the different stability formulations in lateral raise task. Gray lines show joint moments, and colored lines show reserve actuators moments with the different stability formulations. Compared to the joints’ moments, reserve actuator moments were negligible ensuring movement generated by muscles. Only with the Point model, the elbow flexion reserve actuator moment was unrealistically higher than the joint moment.

**Fig S6.**
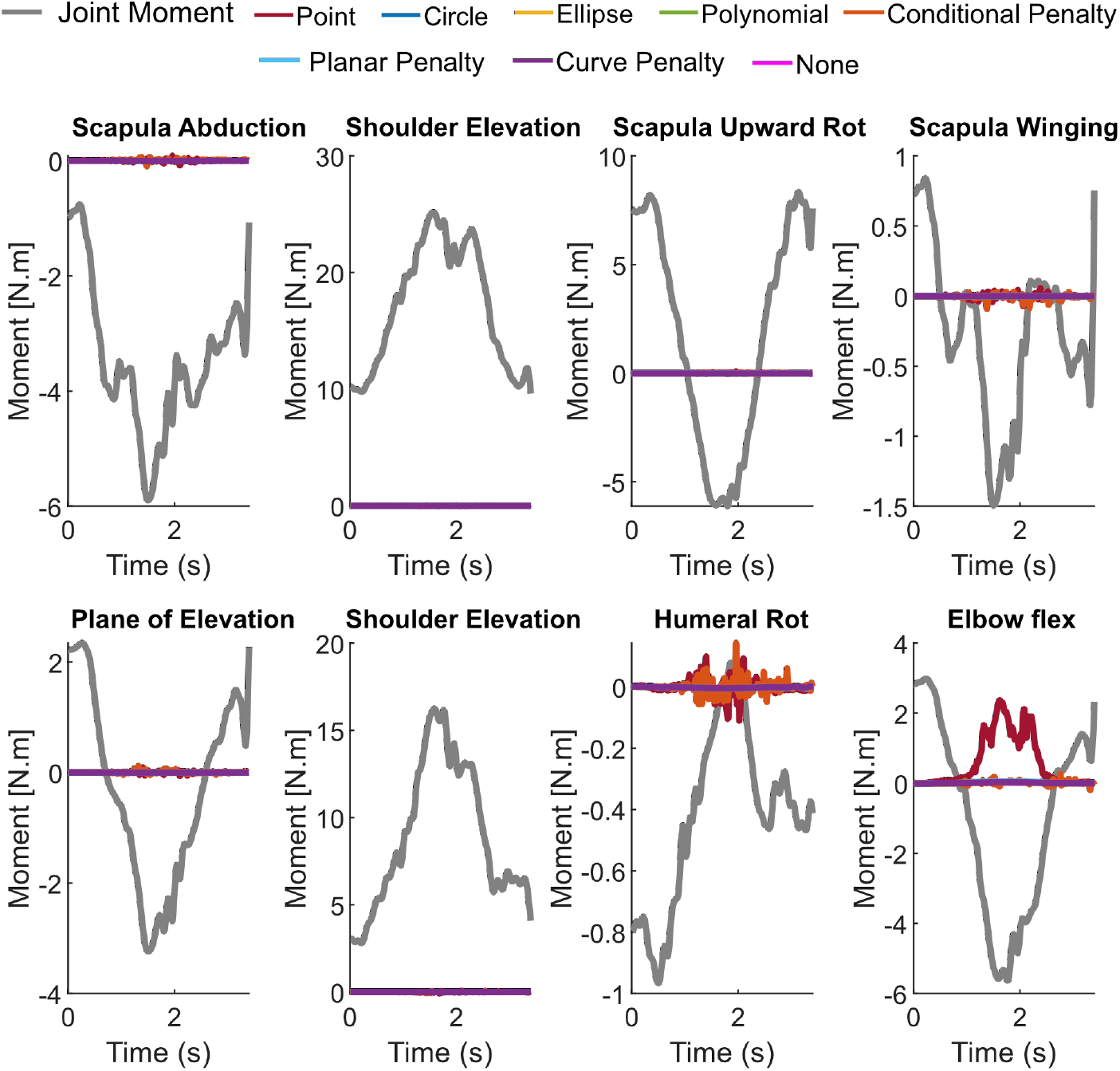
Moments of the scapulothoracic joint and the GH joint compared to moments of reserve actuators with the different stability formulations in posterior raise task. Gray lines show joint moments, and colored lines show reserve actuators moments with the different stability formulations. Compared to the joint moments, reserve actuator moments were negligible ensuring movement generated by muscles. Only with the Point model, the elbow flexion reserve actuator moment was considerably high compared to the elbow flexion moment.

**Fig S7.**
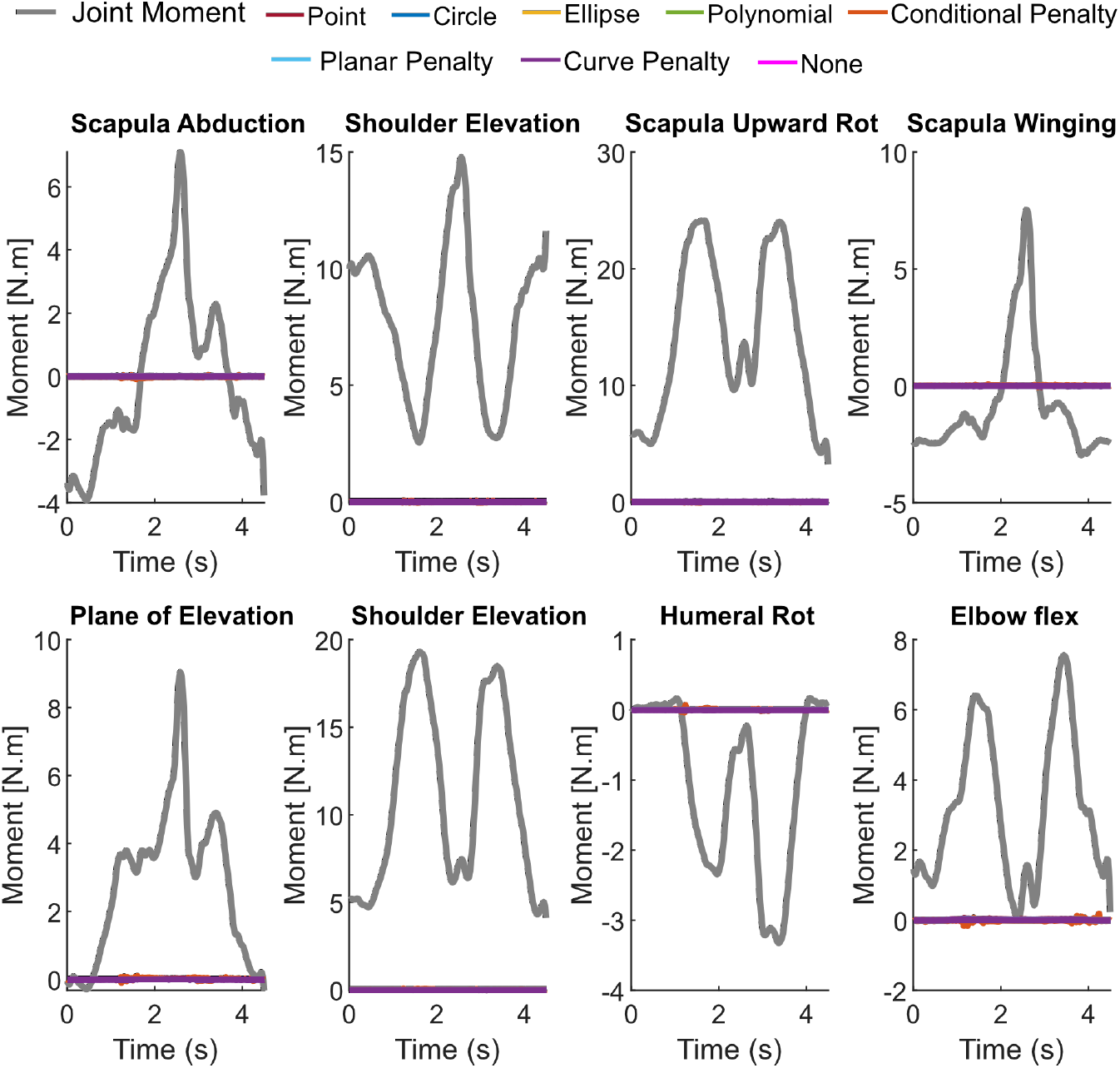
Moments of the scapulothoracic joint and the GH joint compared to moments of reserve actuators with the different stability formulations in Anterior raise task. Gray lines show joint moments, and colored lines show reserve actuators moments with the different stability formulations. Compared to the joint moments, reserve actuator moments were negligible ensuring movement generated by muscles.

## Notes

### Competing Interest Statement

The authors have declared no competing interest.

